# scMontage: Fast and Robust Gene Expression Similarity Search for Massive Single-cell Data

**DOI:** 10.1101/2020.08.30.271395

**Authors:** Tomoya Mori, Naila Shinwari, Wataru Fujibuchi

**Affiliations:** Center for iPS Cell Research and Application (CiRA), Kyoto University, 53 Kawahara-cho, Shogoin, Sakyo-ku, Kyoto 606-8507, Japan; Bioinformatics Center, Institute for Chemical Research, Kyoto University, Gokasho, Uji, Kyoto 611-0011, Japan

**Author notes:** These authors contributed equally to this work (Tomoya Mori, Naila Shinwari). Corresponding author: (Fujibuchi W).

**Keywords:** Human Cell Atlas, Massive single-cell data, Gene expression profile similarity, Cell type analysis, Fisher Z-transformation

## Abstract

Single-cell RNA-seq (scRNA-seq) analysis is widely used to characterize cell types or detect heterogeneity of cell states at much higher resolutions than ever before. Here we introduce scMontage (https://scmontage.stemcellinformatics.org), a gene expression similarity search server dedicated to scRNA-seq data, which can rapidly compare a query with thousands of samples within a few seconds. The scMontage search is based on Spearman’s rank correlation coefficient and its robustness is ensured by introducing Fisher’s Z-transformation and Z-test. Furthermore, search results are linked to a human cell database SHOGoiN (http://shogoin.stemcellinformatics.org), which enable users to fast access to additional cell-type specific information. The scMontage is available not only as a web server but also as a stand-alone application for user’s own data, and thus it enhances the reliability and throughput of cell analysis and helps users gain new insights into their research.

## Introduction

Technology for single-cell analysis has evolved to reveal cell profiles at much higher resolutions than ever before. As an example, several studies have demonstrated that the computational analysis of single-cell RNA-seq (scRNA-seq) data can discover novel cells or cell subtypes. The recently launched Human Cell Atlas (HCA) project [1] is expected to further accelerate the production of single-cell data on an extraordinary scale. These unprecedented massive-scale data will be available to the public through International Nucleotide Sequence Database Collaboration (INSDC) sites such as the Gene Expression Omnibus (GEO) [2] and the Sequence Read Archive (SRA) [3]. Thus, data mining by very fast gene expression profile similarity searches has become increasingly important in terms of screening, clustering, and finding cells.

The concept of similarity searches for gene expression profiles was proposed nearly 20 years ago [4]. CellMontage [6] is the first practical and large-scale implementation that provides users quick searches against a large-scale microarray database for similar gene expression profiles based on Spearman’s rank correlation.

Here, we propose scMontage, a renovated gene expression similarity search server, which is developed for analyzing massive-scale scRNA-seq data, based on the SHOGoiN human cell type database (http://shogoin.stemcellinformatics.org) with statistically robust Fisher’s Z-transformed correlation coefficient. Currently, the scMontage server provides human and mouse scRNA-seq data and allows users to quickly access cell-type-specific biological information, such as cell taxonomy, lineage map, cell marker, and so on. The scMontage enhances the throughput and reliability of single-cell analysis and helps users gain new insights into massive scRNA-seq data.

## Results

A profile search in scMontage can be implemented by selecting a database and inputting a query profile. After selecting the database by specifying the organism and the platform, the user can limit the genes for calculation to particular types according to Gene Ontology [7]. As a query, it is possible to either upload a gene expression profile or directly paste gene expression data in CM format.

**Figure 1** shows an example screen shot when human pancreatic alpha cell (GEO id: GSM1901473) is queried to the database, where ‘H. sapiens’, ‘HiS-eq2000/2500’, and ‘MF:transcription factor activity, protein binding’ are selected. The results show that the first hit is the query itself, as expected, and the top hits come from the pancreas alpha cells (**Table 1**, Table S1). The description column contains SHOGoiN Cell IDs (in parentheses) from which a user can access integrated cell type information by the SHOGoiN database. Similarly, when human pancreatic islet cell (GEO id: GSM1901455) is queried to the database under the same database setting as the previous search, the pancreatic islet cell sample is found in the top hits with high statistical significance though less number of pancreatic islet cell samples are contained in the database than the other pancreatic cells (Table S2). The reliability of the scMontage search is not limited to human cell samples. Table S3 shows the search result when mouse Reg4-positive intestinal cell is queried to the database of “M. musculus, SINGLECELL: all” with “MF:transcription factor activity protein binding” genes. The top hit is the Reg4-positive intestinal cell and most of the top hits are small intestinal cells. Therefore, the scMontage search is robust not only for cell types but also for species.

**Figure 1.**
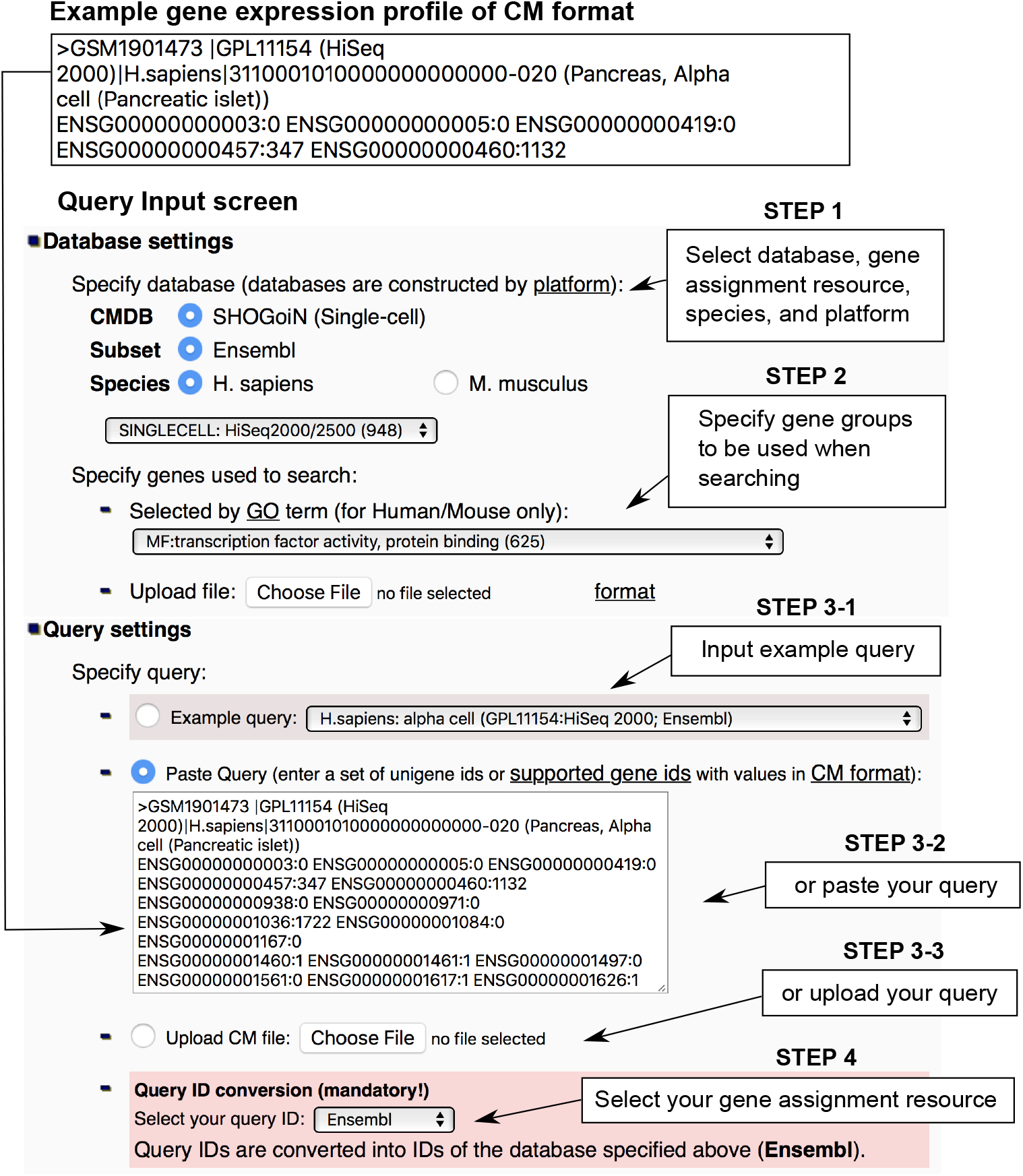
Example of CM format and input screen of scMontage. This example screen shot shows the case that human pancreatic alpha cell is queried to the database. ‘H. sapiens’, ‘HiSeq2000/2500’, and ‘MF:transcription factor activity, protein binding’ are selected as database settings. The search results are shown in Table 1.

**Table 1.** Example search result.

| Sample id | Description | Correlation (P-value of Fisher's Z-transformed rank correlation coefficient) |
| --- | --- | --- |
| GSM1901473 | Pancreas, Alpha cell<br>(3110001010000000000000-020) | 1.0<br>(0.0) |
| GSM1901487 | Pancreas, Alpha cell<br>(3110001010000000000000-020) | 0.56<br>(3.6e-13) |
| GSM1901493 | Pancreas, Alpha cell<br>(3110001010000000000000-020) | 0.54<br>(7.2e-12) |
| GSM1901488 | Pancreas, Alpha cell<br>(3110001010000000000000-020) | 0.53<br>(1.9e-10) |
| GSM1901458 | Pancreas, PP cell<br>(3110001010000000000000-212) | 0.53<br>(3.7e-10) |
| GSM1901497 | Pancreas, Alpha cell<br>(3110001010000000000000-020) | 0.52<br>(1.1e-09) |
| GSM1901464 | Pancreas, Duct cell<br>(3110002050000000000000-090) | 0.52<br>(1.6e-09) |

*Note:* Search result when human pancreatic alpha cell (GEO id: GSM1901473) is queried. Numbers in “Description” indicate SHOGoiN Cell IDs.

In addition, **Figure 2** shows a comparison of statistical evaluation between CellMontage and scMontage for a mouse lung cell sample (GEO id: GSM1271921) under the database setting of “SINGLECELL: all” and “MF:transcription factor activity, protein binding”. The histograms indicate the distributions of the Spearman’s rank correlation coefficient *r*, the t-statistic 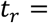 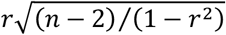, the Z-transformed sample correlation coefficient *z*_*r*_, and the standardized Z-transformed sample correlation coefficient *z*, respectively. In CellMontage, the distribution of *t*_*r*_ does not follow t-distribution when the population correlation coefficient between query and database profiles is non-zero. In scMontage, however, the distribution of *z*_*r*_ approximately follows the normal distribution whose mean is *z*_*ρ*_ = 0.42. Consequently, the standardized Z-transformed sample correlation coefficient *z* follows the standard normal distribution.

**Figure 2.**
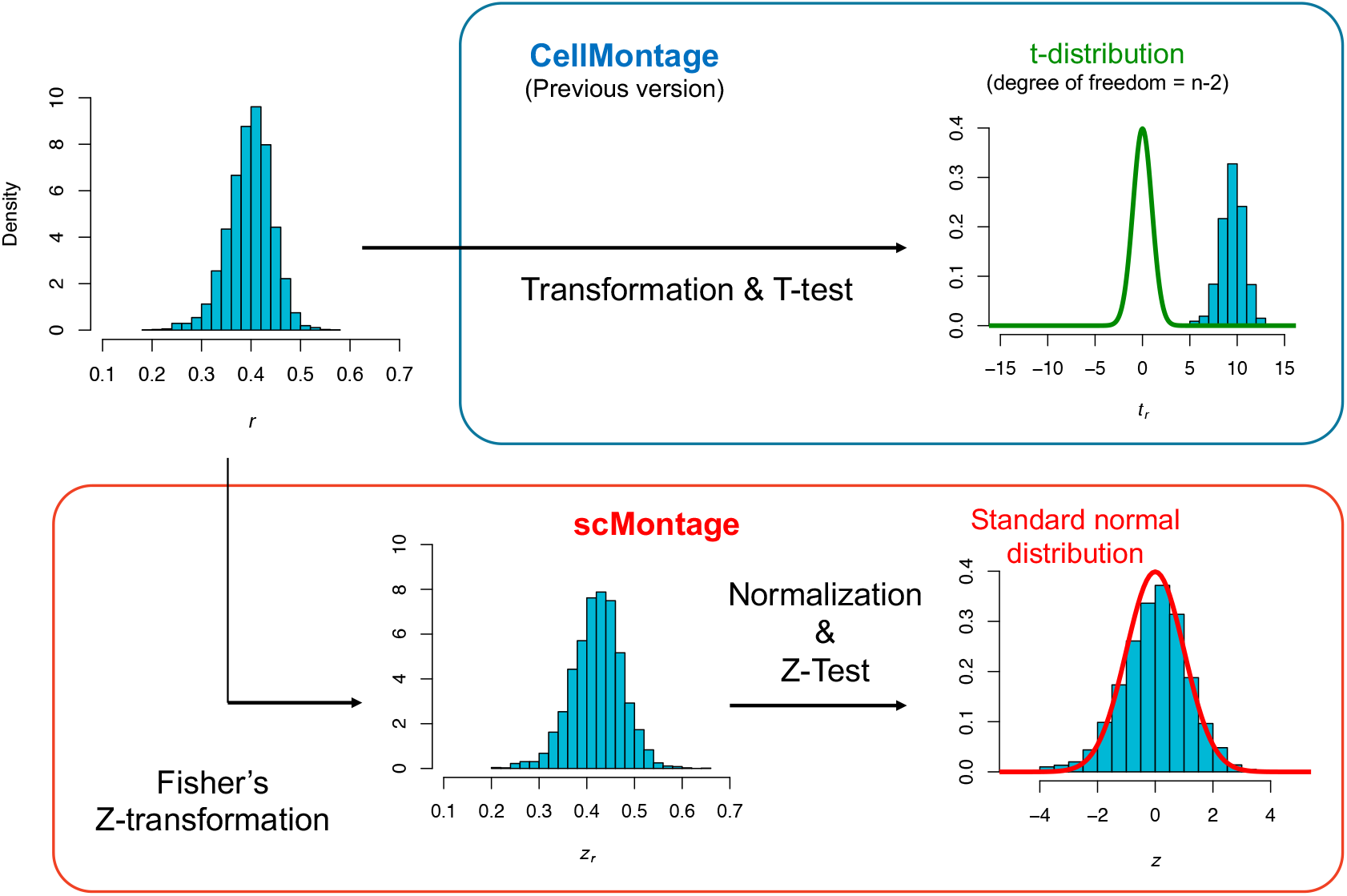
Statistical evaluations of search results from CellMontage and scMontage approaches. The histograms indicate the distributions of the sample correlation coefficient *z*_*r*_, the t-statistic *t*_*r*_ the Z-transformed sample correlation coefficient *z*_*r*_, and the standardized Z-transformed sample correlation coefficient *z* when a mouse lung cell sample (GSM1271921) is queried to the database under the database setting of “SINGLECELL: all” and “MF:transcription factor activity, protein binding”.

## Discussion

We developed scMontage that can be used for the validation and functional prediction of unknown cell types obtained from tissues or derived from stem cells at the single-cell level. The scMontage also provides quick access to additional information of various cell types in the SHOGoiN database from the search results. It is highly expected that a vast amount of single-cell gene expression profiles will be produced from the HCA projects or other research groups in the future. Therefore, scMontage will become an important tool for providing a very fast and powerful environment that can accelerate massive single-cell data analysis by extracting information on gene expression similarity relationships between known and unknown as well as within known/unknown cell types.

## Materials and methods

The scMontage basically runs on Spearman's rank correlation coefficient as a similarity metric of gene expression profiles using a very fast algorithm, RaPiDS [8], for vast calculation, which enables a linear time search with a small constant for the size of the database. As a result, scMontage can compare a query with tens of thousands of samples in the database within a minute. The Spearman’s rank correlation coefficient *r* between two rank numbers is defined as

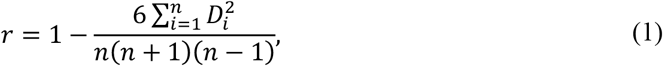

where *D*_*i*_ and *n* indicate the rank difference between gene *i* and the number of genes to be used for calculation, respectively. As the output of scMontage, profiles with the highest similarity to the query are ranked by their statistical significance on the basis of the Fisher's Z-transformation of the rank correlation coefficient, which is drastically improved from the CellMontage approach. The distribution of Fisher’s Z-transformed sample correlation coefficient *z*_*r*_ approximately follows the normal distribution with a mean *z*_*ρ*_ and a standard deviation 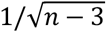 regardless of the size of *n*, where *z*_*ρ*_ is approximated as the mean of *z*_*r*_ when it appears in standardization as the following equations:

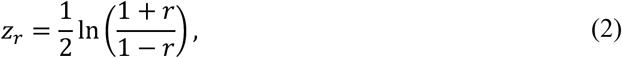

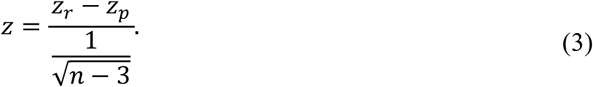

Thus, scMontage can correct the significance that the population correlation coefficient between query and database profiles is non-zero, which often occurs due to common cell properties such as cell cycle states observed at single-cell level regardless of cell types.

The scMontage server currently provides 5,035 single-cell transcriptome data (1,424 human and 3,611 mouse cell samples on 23 August 2018) whose cell types are available by original submitters. Raw sequence data are acquired from SRA, and their read counts are computed by mapping them to human/mouse reference genome sequences downloaded from Ensembl [9] using Bowitie2 [10] and counting the mapped reads by HTSeq [11].

Furthermore, scMontage results are linked to the SHOGoiN database, a repository for accumulating, integrating, and providing cell information of human and other model organisms. This allows users fast access to additional cell-type-specific information, such as cell taxonomy, lineage map, cell marker, DNA methylation, and morphological image.

## Authors’ contributions

WF conceptualized and designed the study. TM, NS, and WF developed the server and drafted the paper. All authors have read and approved the final manuscript.

## Acknowledgements

This work was partially supported by the Core Center for iPS Cell Research, Research Center Network for Realization of Regenerative Medicine, Japan Agency for Medical Research and Development, Grant-in-Aid for Scientific Research on Innovative Areas, The Ministry of Education, Culture, Sports, Science and Technology, and the iPS Cell Research Fund. The authors deeply appreciate Dr. Peter Karagiannis for kindly reviewing the manuscript.

## Supplementary material

**Supplementary Table S1 Search result when human pancreatic alpha cell (GEO id: GSM1901473) is queried**

**Supplementary Table S2 Search result when human pancreatic islet cell (GEO id: GSM1901455) is queried**

**Supplementary Table S3 Search result when mouse Reg4-positive intestinal cell (GEO id: GSM1524296) is queried**

